# Evolutionary Reconstruction of Hormone-bHLH Regulatory Networks in Solanaceae: Phylogenomic Insights from PSTVd-Tomato Interactions

**DOI:** 10.1101/2025.03.14.643413

**Authors:** Katia Aviña-Padilla, Octavio Zambada-Moreno, Michelle Bustamante Castillo, Manuel A. Barrios-Izás, Maribel Hernández-Rosales

## Abstract

The evolutionary reconstruction of transcriptional regulatory networks play a crucial role in plant adaptation to biotic stress. In Solanaceae, basic helix-loop-helix (bHLH) transcription factors (TFs) regulate hormone signaling pathways that balance growth and defense responses. In this study, we investigated the phylogenomic evolution of bHLH-mediated hormone networks in response to Potato Spindle Tuber Viroid (PSTVd) infection in tomato (*Solanum lycopersicum*) and its wild relatives. Among them, the JA-associated bHLH *Solyc08g076930*, IA3/MYC2 exhibits strong evolutionary conservation across Solanaceae, whereas the auxin regulator *Solyc03g113560* miP-bHLH-ARF8 is a tomato-specific gene raising from an ancestral duplication, indicating neofunctionalization. Under PSTVd infection, we observe extensive regulatory rewiring, with bHLH hubs assuming dominant control over target genes typically regulated by multiple TFs in healthy conditions. The JA regulon undergoes the most pronounced shift, with *Solyc08g076930* directly regulating 91 genes, compared to shared regulation with over a thousand TFs in healthy plants. This rewiring reflects the adaptive plasticity of bHLH TFs in coordinating stress responses through hormonal crosstalk. These findings reveal key evolutionary pressures shaping Solanaceae stress responses, offering insights for improving viroid tolerance and crop resilience.

## 1. Introduction

The evolutionary history of transcriptional regulatory networks offers critical insights into plant adaptation to biotic stress [1]. Within these networks, basic Helix-Loop-Helix (bHLH) transcription factors (TFs) form a highly conserved family central to stress responses, hormone signaling, and development [2–4]. By binding E-box motifs (CANNTG), bHLH TFs modulate gene expression, driving key metabolic and adaptive processes. They are classified into six groups: Class I (bHLH-b), including MYC2, a core jasmonic acid signaling regulator [5]; Class II (bHLH-A), featuring PIFs involved in light responses [6]; Class III (bHLH-PAS), which contains PAS domains for environmental sensing [7]; Class IV (bHLH-B), functioning as passive repressors [8]; Class V, comprising factors like BEE proteins linked to brassinosteroid signaling [9]; and Class VI (MYC-like), such as TT8, which regulates secondary metabolism and flavonoid biosynthesis [10]. Despite their recognized roles, their evolutionary trajectories under pathogen-induced stress remain poorly understood [4,11]. Our prior research demonstrated their pivotal role in tomato transcriptional reprogramming during Potato Spindle Tuber Viroid (PSTVd) infection [20–21]. PSTVd, a subviral RNA pathogen, disrupts host gene networks, causing severe developmental abnormalities and economic losses in major crops [15]. Unlike viruses, viroids lack coding potential, relying exclusively on the host’s transcriptional machinery for replication and systemic movement [16]. In this context, the Solanaceae family represents a relevant model for investigating bHLH-mediated regulatory responses to viroid infection due to its agricultural significance and the varying degrees of viroid tolerance among its members. Notably, wild species such as *Solanum pennellii*, *S. pimpinellifolium*, and *Capsicum annuum* var. *glabriusculum* exhibit greater viroid tolerance compared to their domesticated counterparts, including *S. lycopersicum*, *S. lycopersicum* var. *cerasiforme*, and *S. tuberosum* [13–14,17–19]. This differential response provides a framework for understanding the molecular mechanisms underlying viroid tolerance and susceptibility in economically important crops. The observed differences in susceptibility suggest that evolutionary pressures have shaped distinct regulatory architectures in viroid hosts. Reconstructing the evolutionary history of hormone-mediated regulation can uncover the molecular basis of stress adaptation, highlighting key regulatory elements associated with resilience or susceptibility. Additionally, exploring the diversification and specialization of these transcription factors across species offers insights into the evolutionary dynamics of plant defense strategies. Complementary, regulatory network analysis provides a systems-level perspective on how PSTVd disrupts co-expression patterns and transcription factor interactions, driving transcriptional reprogramming under viroid stress. Together, these approaches offer a comprehensive understanding of the evolutionary and functional mechanisms shaping viroid responses in natural hosts.

In this study, we investigate bHLH-mediated regulation in plant responses to PSTVd infection, focusing on three TFs associated with hormonal signaling. These include MYC2, a master transcriptional regulator (MTR) of jasmonic acid (JA) pathways, and two bHLH microproteins: one associated with auxin signaling and the other with brassinosteroid pathways. Through the integration of phylogenomic reconstruction, transcriptomic profiling, and network analysis, we uncover how bHLH regulators orchestrate gene expression during viroid stress, highlighting their role in coordinating responses in natural viroid hosts. Our findings offer valuable insights for advancing strategies to enhance disease resistance and stress resilience in Solanaceae crops, with potential applications in breeding and genetic engineering to support sustainable agriculture.

## 2. Materials and methods

### 2.1 Data extraction of bHLH TFs in domesticated and wild tomato

A total of 161 bHLH TFs were identified in the domesticated tomato (*S. lycopersicum*), and 172 in the wild species (*S. pennellii*), serving as the basis for comparative genomic analysis in the evolutionary reconstruction. These proteins were retrieved from PlantTFDB (https://planttfdb.gao-lab.org), an experimentally curated database for plant TFs, ensuring high-confidence annotation and classification. All codes used for this analysis are available at: https://github.com/ozambadam/Hormone-Orthologs-Analysis-Tomato/tree/main. The complete methodology is described in **Fig. A1**.

### 2.2. Host selection and data acquisition

To assess viroid tolerance across Solanaceae species, we selected three wild and four domesticated hosts:*S. pennellii, S. pimpinellifolium, C. annuum var. glabriusculum, and S. lycopersicum, S. lycopersicum var. cerasiforme, C. annuum, S. tuberosum*), categorized by tolerance level and associated symptoms as shown in **Appendix Table 1**. Complete proteomes for these species were downloaded from the Solgenomics Network DB (https://www.solgenomics.net) in FASTA format.

### 2.3 Evolutionary Reconstruction Across Domesticated and Wild Species

To explore the evolutionary conservation and functional divergence of bHLH transcription factors (TFs) across seven Solanaceae species, we employed REvolutionH-tl, a computational framework for gene family evolution analysis [22]. This method reconstructs event-labeled gene trees, where internal nodes represent duplication or speciation events. These trees are reconciled with the species tree to pinpoint lineage-specific duplications and gene losses, providing insights into the evolutionary trajectories of bHLH TFs. The species tree inference follows a multi-step process. First, sequence alignments across species are performed to identify homologous sequences, generating key alignment statistics such as bit-score and e-value. The most evolutionarily related genes are identified as best hits, and orthogroups are defined as sets of genes descending from a common ancestor. Based on this classification, gene trees are reconstructed, and orthology is assigned to conserved genes. Consensus speciation signals across gene trees inform the species tree construction. The reconciliation step integrates gene trees into the species tree, allowing the identification of gene duplications, losses, and lineage-specific expansions.

### 2.4. Transcriptomic data acquisition and Integration for the Gene Regulatory Network (GRN)

Microarray datasets of root (GSE111736) and leaf (GSE106912) transcriptomes from *S. lycopersicum* under mild and severe PSTVd infection were retrieved from NCBI GEO (https://www.ncbi.nlm.nih.gov/geo). The datasets (26 root, 27 leaf samples) were merged into a unified expression matrix [23–24]. Gene duplicates were averaged, and data were normalized using the RMA function from the affy package. To enhance network building, we merged the samples to obtain a more comprehensive representation of gene expression patterns across different infection conditions. The final dataset comprised 53 samples: 18 control (C), 35 PSTVd infected samples, ensuring balanced tissue representation.

### 2.5 Hormone PSTVd-Tomato Gene Regulatory Network (GRN) Construction

To investigate *S. lycopersicum* transcriptional regulation in response to PSTVd infection, GRN was constructed using the Corto R package. Processed and normalized transcriptomic data enabled the identification of bHLH TFs as MTRs. For this analysis, we used an integrated expression matrix and the PlantTFDB tomato transcription factor list. Corto employs Spearman correlation to identify gene associations and applies the Data Processing Inequality (DPI) to eliminate indirect interactions, with bootstrapping to ensure robustness. The GRN network included a total of 53 samples and 8080 features. Spearman correlation thresholds of 0.65514 were applied, alongside a p-value threshold of 1 × 10 was applied to filter significant edges, and 100 bootstraps were performed to ensure robustness. From these global GRN, subnetworks focusing on bHLH-TFs and their interactors were extracted, enabling detailed analysis of tissue-specific regulatory dynamics under PSTVd-induced stress.

### 2.6 Master Transcriptional Regulator (MTR) and microprotein (miP) Identification

Given that bHLH TFs exert their biological roles by regulating target genes, we infer their functional involvement through the analysis of their regulons. To achieve this, functional enrichment analysis was performed using the gProfiler R package to identify biological processes. Significantly enriched terms were determined using a false discovery rate (FDR) threshold of <0.05. miP assignment was assessed using the miPFinder2 tool [25]. PPI data from STRING DB (https://string-db.org) refined regulatory associations, uncovering potential mechanisms underlying transcriptional and posttranscriptional reprogramming. Finally, bHLH regulatory subnetworks linked to a JA MTR, and two miPs related to auxin, and brassinosteroid signaling pathways were extracted to explore interactions in the viroid response.

### 2.7 Alternative Regulator Identification in Domesticated and Wild Tomato Networks

Custom scripts in Bash and R analyzed regulatory networks of domesticated and wild type species and identified conserved and divergent regulatory roles. Using the *S. lycopersicum* PlantTFDB regulatory map (https://planttfdb.gao-lab.org), we identified conserved bHLH elements and rewiring events in healthy vs disease states. Using *S. pennellii’s* PlantTFDB regulatory map, we also identified conserved bHLH genes in its regulatory network. For divergent bHLH TFs, functional annotation revealed lineage-specific regulatory shifts and compensatory mechanisms.

## 3. Results

### 3.1 Evolutionary Reconstruction of bHLH Transcription Factors in Solanaceae

The evolutionary history of TFs plays a critical role in shaping GRNs, particularly in response to selective pressures such as domestication and environmental adaptation. In this study, we investigate the evolutionary trajectories of bHLH TFs across Solanaceae, with a focus on domesticated tomato (*S. lycopersicum*) and its wild relative (*S. pennellii)*. Through a phylogenomic approach integrating gene gain, loss, and duplication events, we reconstruct the evolutionary history of this gene family and explore its implications for transcriptional regulation, **Fig. 1**.

**Fig. 1.**
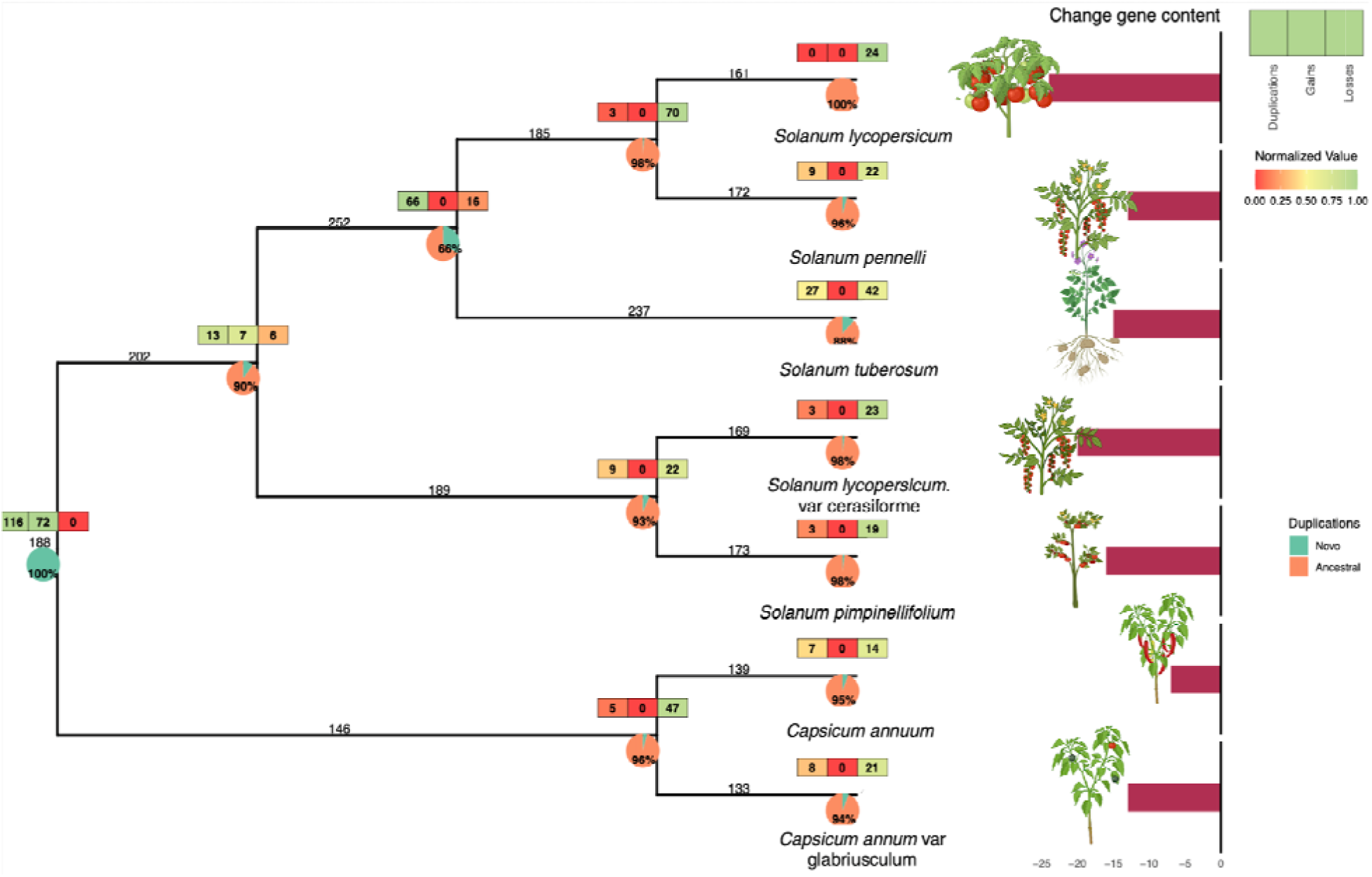
Evolutionary reconstruction of bHLH TFs in domesticated and wild tomato. This phylogenomics analysis illustrates the evolutionary trajectory of bHLH genes across domesticated and wild Solanaceae species. Gains (green), losses (yellow), and duplications (red) are mapped onto branches, with pie charts indicating the proportion of novel (green) and ancestral (orange) duplications at each evolutionary node. The accompanying bar plot quantifies gene content changes across lineages.

The *S. lycopersicum* genome contains 161 bHLH TFs, with 159 conserved and 2 species-specific. *S. pennellii*, its closest wild relative, has 172 bHLH,, also with 2 species-specific genes but 9 lineage-specific duplications leading to co-orthologs. The ancestral Solanaceae genome contained 188 bHLH TFs, with substantial losses in Capsicum and expansions in Solanum. For instance, *S. tuberosum* (potato) has 237 bHLH genes, many retained from an ancestral expansion, but later lost in *S. lycopersicum* and *S. pennellii*. Notably, all species show a net reduction in gene content compared to their last common ancestor, likely reflecting functional specialization. Moreover, our results highlight lineage-specific expansions and contractions shaping bHLH evolution. In the common Solanum ancestor, 13 duplications, 7 gene gains, and 6 losses marked the start of regulatory divergence. This intensified in the ancestor of *S. lycopersicum, S. pennellii,* and *S. tuberosum*, with 66 duplications and 16 losses, followed by 3 additional duplications and 70 losses in the *S. lycopersicum*-*S. pennellii* lineage. Domesticated tomato lost 24 bHLH genes, likely due to selection during domestication, while *S. pennellii* lost 22 genes but gained 9 through duplications, reflecting a parallel yet distinct evolutionary path.While closely related species (*S. pimpinellifolium*, *S. cerasiforme*) maintain a more conserved bHLH repertoire, *S. tuberosum* underwent extensive gene turnover, with 27 species-specific duplications and 42 losses, suggesting adaptation to unique developmental and stress-response pathways. Similarly, Capsicum species experienced significant ancestral losses (47 genes) with only 3 duplications, indicating distinct regulatory adaptations.

The reconstruction of all the BHLH genes found in domesticated (*S. lycopersicum*) and wild (*S. pennellii*) tomato species reveals that *S. lycopersicum* possesses one differential bHLH TF *Solyc10g079660.* Interestingly, this gene is also found in other domesticated species belonging to the Solanum spp such as *S. lycopersicum* var cerasiforme and *S. tuberosum.* Using Protein-Protein Interactions from STRING (https://string-db.org/) data we found its functional enrichment that suggest that the identified interacting genes play a crucial role in intracellular transport mechanisms, particularly in nucleocytoplasmic trafficking, functions that may be conserved across domesticated species. Moreover, our analysis identifies two unique bHLH TFs exclusive to *S. lycopersicum*, *Solyc02g067380* and *Solyc06g070910*. Functional enrichment analysis of their interactomes highlights significant roles in cytoskeletal dynamics and hormonal signaling. Specifically, they are associated with *microtubule depolymerization* and *microtubule-based movement*, processes essential for intracellular transport and cell division. Additionally, their involvement in *steroid hormone* and *brassinosteroid-mediated signaling pathways* suggests roles in growth regulation, while enrichment in *cellular response to lipid*, points to lipid-mediated signaling functions. While the precise roles of these genes in pathogen susceptibility have not been fully elucidated, hormonal pathways are known to significantly influence plant defense mechanisms.

### 3.2 Hormonal Crosstalk and Evolutionary Divergence in PSTVd-Infected Tomato: Insights from Network and Ortholog Analysis

The regulatory network analysis of *S. lycopersicum* response to PSTVd infection reveals a hierarchical model integrating auxin, jasmonic acid, and brassinosteroid signaling (**Fig. 2a**), [20]. *Solyc08g076930* (IA3/MYC2, MTR) emerges as a central JA regulator, linking iron homeostasis, hydrocarbon biosynthesis, and transcriptional adaptation. Its dual role in PPI interactions and co-expression networks suggests that IA3/MYC2 fine-tunes growth, defense, and metabolic transitions under viroid induced stress conditions. Meanwhile, *Solyc03g113560* (miP-bHLH-ARF8 modulator) acts as a regulatory hub, integrating auxin signaling through PPI interactions, while displaying co-expression enrichment in methionine biosynthesis, amino acid metabolism, and ribosomal processes, indicating a role in translational control during stress. Similarly, *Solyc05g007210* (miP-bHLH Brassinosteroid-Linked modulator) bridges brassinosteroid signaling with lipid metabolism and energy regulation, interacting via PPI with ILI1-binding factor and exhibiting co-expression enrichment in fatty acid desaturation, pyruvate kinase activity, and ion transport, suggesting a function in membrane remodeling and metabolic homeostasis.

**Fig. 2.**
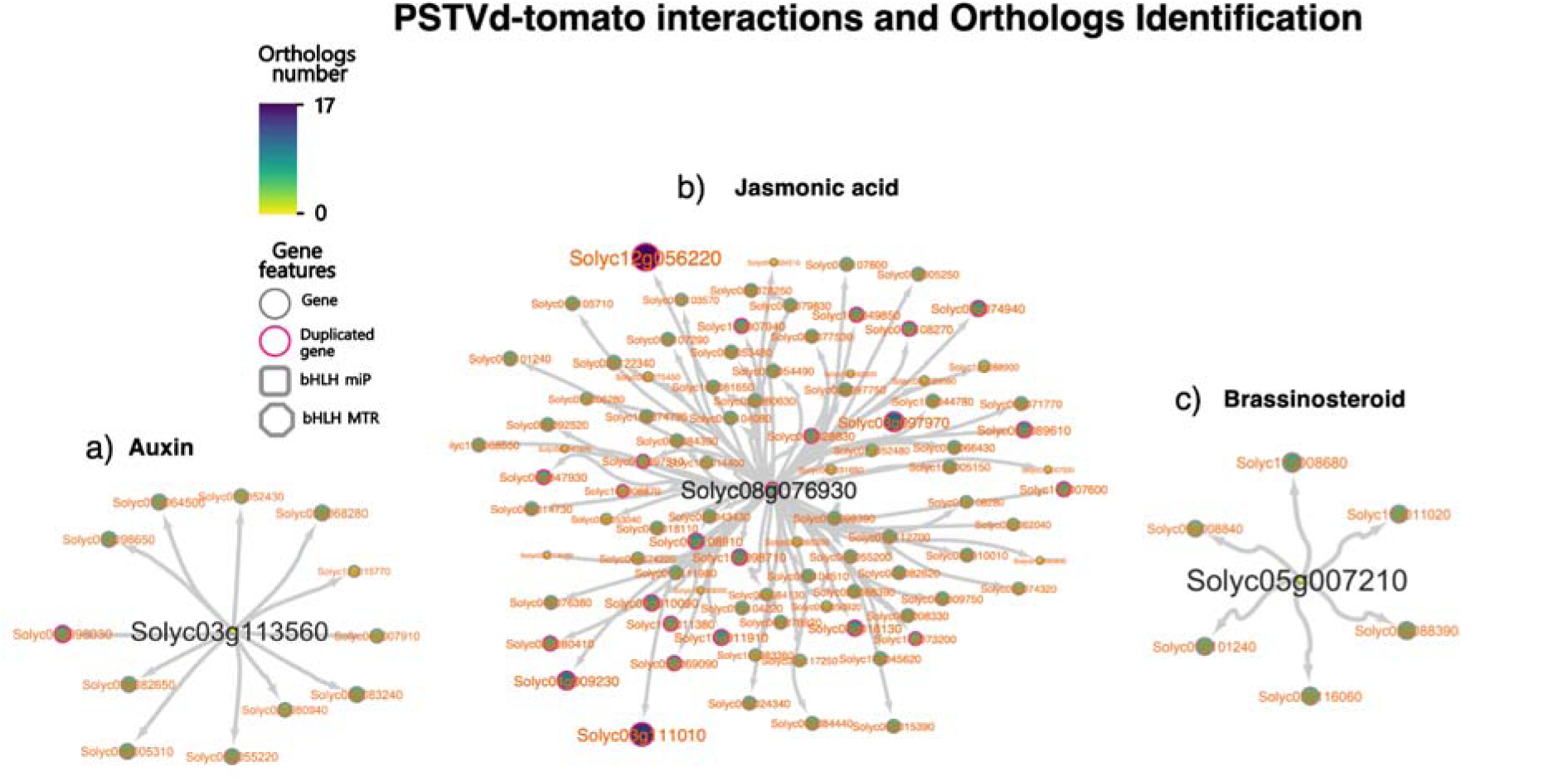
PSTVd Regulatory Network and Orthologs in Tomato. The networks depict the regulatory modules associated with (a) auxin (*Solyc03g113560*), (b) jasmonic acid (*Solyc08g076930*), and (c) brassinosteroid (*Solyc05g007210*) signaling pathways. Nodes represent genes, with duplicated genes outlined in pink. Gray squares indicate bHLH miPs, while hexagons denote bHLH master regulators (MTR). The color gradient represents the number of orthologs identified across Solanaceae species, ranging from 0 (yellow) to 17 (purple). This analysis highlights key regulatory hubs modulating transcriptional responses under PSTVd infection.

Ortholog inference further highlights differential conservation among these pathways. IA3/MYC2 regulon exhibit the highest evolutionary reinforcement, suggesting strong selection pressure on JA-mediated regulation (**Fig. 3**). In contrast, miP-auxin regulator shows more restricted retention, reflecting a tighter evolutionary constraint. While miP-Brassinosteroid-linked modulator components maintain conservation, they also exhibit signs of adaptive rewiring across Solanaceae hosts. These findings suggest that signal-metabolic regulatory networks operate under distinct evolutionary constraints, with JA and brassinosteroid TFs regulons showing strong conservation, while auxin exhibit species-specific divergence, shaping stress-adaptive transcriptional responses in *S. lycopersicum*.

**Fig. 3.**
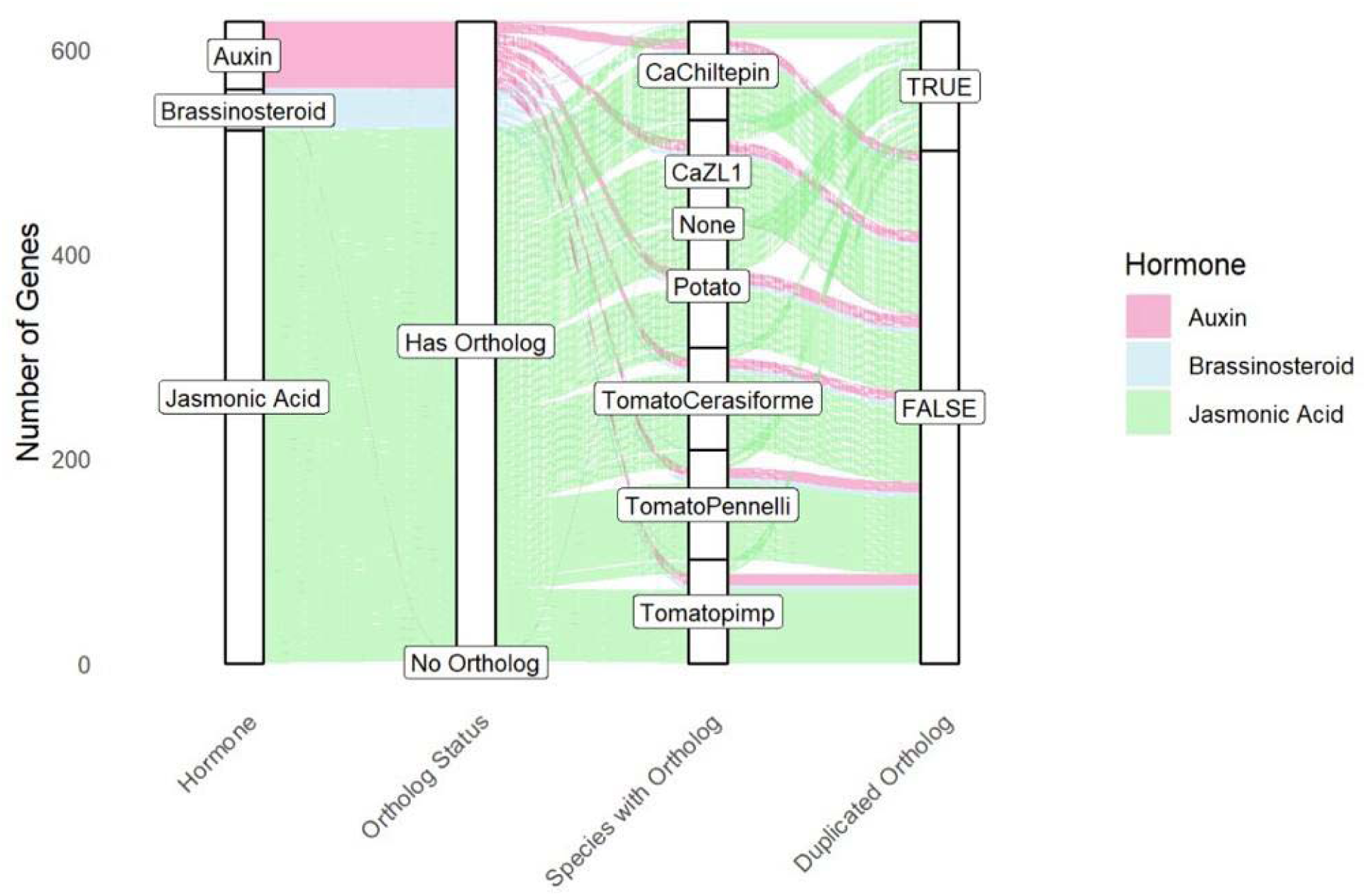
Sankey diagram showing ortholog distribution for genes involved in bHLH-ARF8 (pink), bHLH-brassinosteroid (blue), and IA3/MYC2 (green) regulons across Solanaceae species. The plot displays the number of genes by hormone pathway (left), their ortholog status (middle-left), species with identified orthologs (middleright), and duplication status (right, TRUE if the gene has more than one co-ortholog in a species, FALSE if there are none). Genes related to JA signaling are the most prevalent and widely conserved, while auxin and brassinosteroid genes show more lineage-specific patterns, especially in Solanum species.

#### 3.2.1. miP bHLH-ARF8 regulon: Tomato-Specific Genes and Functional Divergence

The reconstruction of the gene family containing Solyc03g113560, a 103-amino acid bHLH microprotein (miP) with ARF-8 regulatory activity, reveals that despite belonging to a large gene family, it has no orthologs in other species. This gene arose through a duplication event, making it a paralog rather than a direct ortholog to related genes in other species. Its divergence is likely due to neofunctionalization, where it acquired a new function essential for domestication. This bHLH functions as a key hub within the auxin regulatory network during viroid infection (**Fig. 2b**). Its absence in other species confirms it as a tomato-specific gene, suggesting functional divergence in auxin regulation (**Fig. 4a**).

**Fig. 4.**
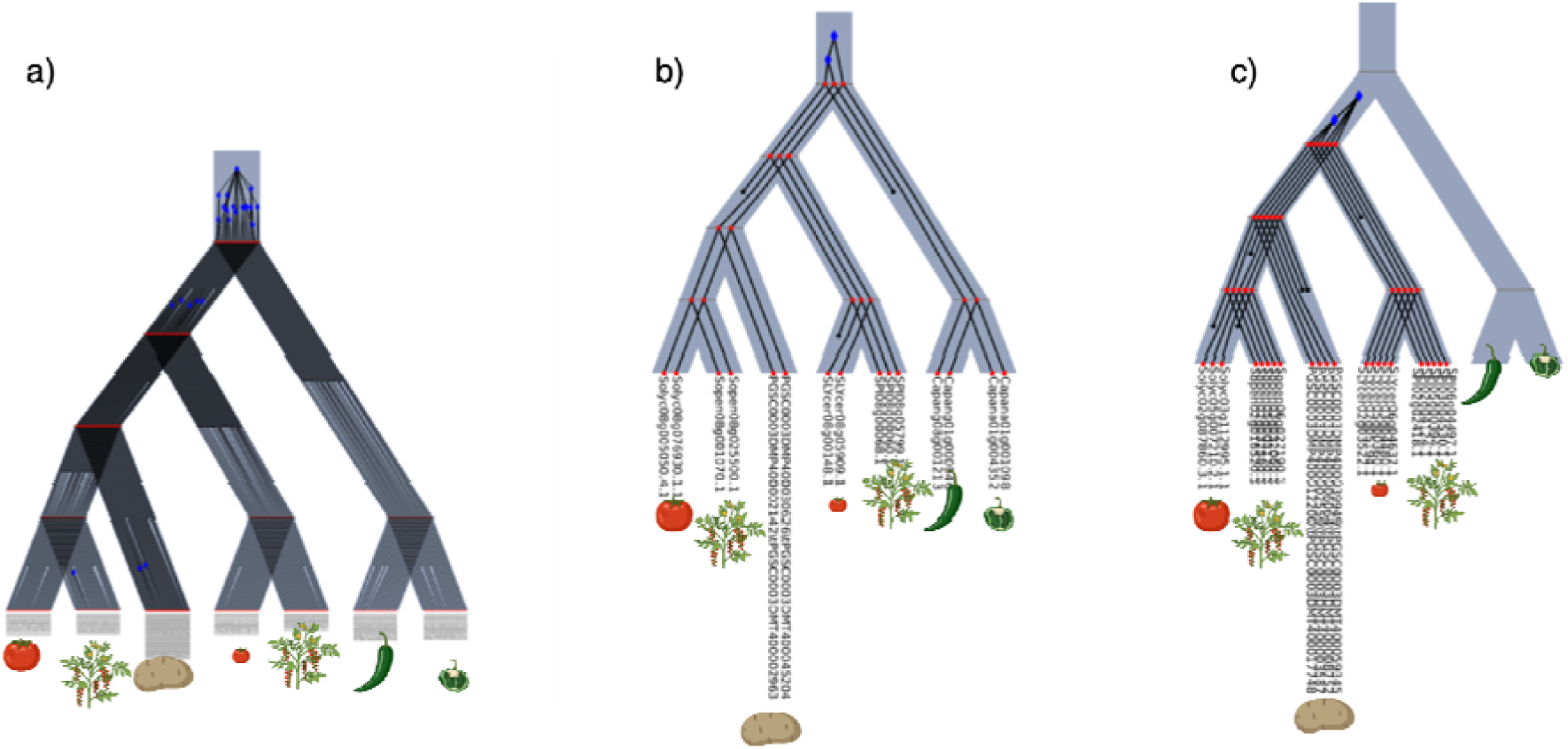
Phylogenetic trees of the a) bHLH-ARF8 (miP-auxin), b) IA3/MYC2 (jasmonate), and c) bHLH-brassinosteroid linked (miP-brassinosteroid) orthogroup. Evolutionary reconstruction of the bHLH-hormone regulator families. This figure shows the reconciliation of gene trees with the species tree for a gene family (orthogroup), emphasizing the evolutionary trajectory of the bHLH hormone TF. Blue nodes represent inferred gene duplication events, black nodes represent gene losses, while red internal nodes indicate ancestral divergence points. Leaves correspond to orthologs from different species.

In the gene family, ortholog retention varies across species, with *S. pennellii* and *S. cerasiforme* (10), *C.* spp. (11), and *S. tuberosum* (12), indicating lineage-specific selective pressures. In contrast, *Solyc09g098030*, a Cytochrome P450, exhibits strong evolutionary conservation across wild and domesticated Solanaceae, with a duplication event identified in wild pepper, suggesting lineage-specific expansion. Our findings indicate its involvement potentially contributing to stress responses and developmental plasticity. This aligns with previous reports highlighting the role of cytochrome P450 enzymes in JA signaling pathways, where JA-Ile levels are regulated through catabolism mediated by CYP family hydroxylases [29]. Additionally, JAZ proteins, acting as metabolic hubs, coordinate crosstalk with other phytohormone signaling pathways, integrating genome-wide responses. Conservation in domesticated tomato and *Capsicum annuum* highlights its broader evolutionary significance, while its absence in *S. pennellii* suggests lineage-specific loss or significant sequence divergence.

#### 3.2.2 IA3/MYC2, MTR regulon: Highly Conserved Across Solanaceae

The bHLH TF *Solyc08g076930* (*IA3/MYC2*), a key regulator of the JA pathway [30], exhibits a highly orthologous conservancy across Solanaceae, **Fig. 4b**. This broad conservation highlights its functional relevance in stress adaptation. There are 113 duplicated orthologs associated with its regulon, **Fig. 3**. The distribution of duplicated genes in the JA pathway reveals notable differences among species. The highest number of duplications is observed in potato (44 duplications), which may be influenced by its tetraploid genome structure, known to retain more gene copies. In *Capsicum* spp, domesticated (18 duplications) and wild (14 duplications) show a considerable presence of duplications, suggesting a broader retention of MYC2-associated genes within the pepper lineage. Among wild tomato relatives, *S. pennellii* (16 duplications) and *S. pimpinellifolium* (12 duplications) exhibit higher numbers of duplications compared to domesticated cherry tomato (*S. lycopersicum var. cerasiforme*, 9 duplications). This pattern may indicate that wild tomato species tend to retain more copies of JA-related genes than domesticated varieties.The highest number of orthologs was identified in *S. tuberosum* (101 orthologs), followed by *S. pennellii* (91 orthologs), while *C.* species and domesticated tomato relatives (*S. pimpinellifolium* and *S. cerasiforme*) displayed slightly lower but consistent ortholog retention (80–83 genes). This pattern suggests that this regulon is strongly conserved, particularly in wild tomato species and *S. tuberosum*, reinforcing its fundamental role in stress resilience mechanisms [30–32]. In addition to MYC2, several bHLH-regulated genes display lineage-specific expansions and duplications, indicating functional diversification within this regulatory network. Functional annotation of duplicated genes reveals potential roles in cellular processes. Notably, *Solyc12g056220*, encoding *SlPIP1.3*, exhibits the highest duplication rates in *S. tuberosum*, *S. cerasiforme*, *S. pennellii*, and *S. pimpinellifolium* (three copies each), suggesting a conserved role in membrane transport regulation under stress conditions. Similarly, *Solyc03g111010,* encoding a *Glyceraldehyde-3-Phosphate Dehydrogenase*, presents the highest duplication levels in *S. pennellii* and *S. pimpenellifolium* (three copies), indicating a possible metabolic adaptation to energy homeostasis during stress. Interestingly, *Solyc03g097970,* annotated as *Erwinia Induced Protein 2*, exhibits lineage-specific duplications, with three copies in *S. tuberosum* and two in *S. cerasiforme*, *S. pennellii*, and *S. pimpenellifolium*. Given its association with plant defense responses, its duplication in Solanum spp. suggests an enhanced role in stress adaptation, reinforcing adaptive mechanisms in wild tomato species. The presence of multiple copies in wild tomato species but not in domesticated tomato (*S. lycopersicum*) suggests that its evolutionary retention is specific to wild relatives, potentially as an adaptive response to environmental pressures. This pattern points out its role as an important component of the Solanaceae stress response network, likely governed by MYC2-mediated pathways, and highlights the impact of natural selection on stress-related gene expansion in wild tomato lineages. *Solyc04g009230,* identified as a Mitosis protein dim1, contains the InterPro domain IPR004123, associated with mRNA splicing factors and thioredoxin-like U5 snRNPs, suggesting a role in RNA processing and post-transcriptional regulation. This function may be relevant to mediated stress responses, where transcriptional control is critical for adaptive regulation. Similarly, *Solyc01g101240,* annotated as an aspartic protease precursor, is likely involved in proteolytic processing, which plays a role in protein turnover, signaling cascades, and stress adaptation. Aspartic proteases have been implicated in hormonal regulation and programmed cell death, processes that are essential for JA-dependent defense mechanisms [33]. These findings suggest that the expansion of bHLH-regulated genes in the JA pathway involves components associated with RNA regulation and proteolysis, reinforcing the idea that post-transcriptional and post-translational modifications contribute to hormonal responses in Solanaceae [30].

#### 3.2.3 miP-Brassinosteroid-Linked Regulon: Dual Network Integration Through Conserved Stress Regulators

*Solyc05g007210*, an 86-amino acid microprotein (miP), emerges as a key regulator within the identified network (**Fig. 2d**). This miP is conserved across *Solanum* species but is absent in *Capsicum*. The *miP*-brassinosteroid-associated regulon displays a conserved evolutionary distribution among Solanaceae species, but with limited ortholog duplication (Fig. 3). The quantitative assessment reveals no duplicated orthologs across all seven species, suggesting restricted expansion within this regulon compared to the MYC2 network. Despite this limited duplication, ortholog retention patterns are consistent across species: *Solanum* spp. each preserve 7 orthologs, indicating stable conservation. Capsicum species show slightly lower retention, maintaining between 5 and 6 orthologs. Interestingly, *Solyc01g101240* (aspartic protease precursor) is present in both the brassinosteroid and MYC2 regulons highlights a potential point of crosstalk between jasmonic acid and brassinosteroid regulators pathways, suggesting a shared role in stress adaptation.

### 3.3 Inference of the potential regulatory mechanisms under biotic stress across host species: Conservation and divergence

After describing the orthology conservation and divergence of the three hormone-associated TFs and their respective regulons across *Solanum* and *Capsicum* species, we focus on proposing regulatory network models aligned with their evolutionary history. To accomplish this, we first identified the original regulon involved in *S. lycopersicum* response to PSTVd. We then inferred the potential regulatory network based on shared orthologs, reconstructing the ancestral regulon from their most recent common ancestor. We examined how regulatory components have been retained or lost across these species. For instance, the Brassinosteroid regulatory network inferred during infection conserves all regulator and target genes in the Solanum lineage but the bHLH regulator *Solyc05g0072*1is not found in the Capsicum lineage, suggesting that this gene emerged in the Solanum lineage and therefore, conservation of the regulatory mechanism can be hypothesized **Fig 5a**.

**Fig. 5.**
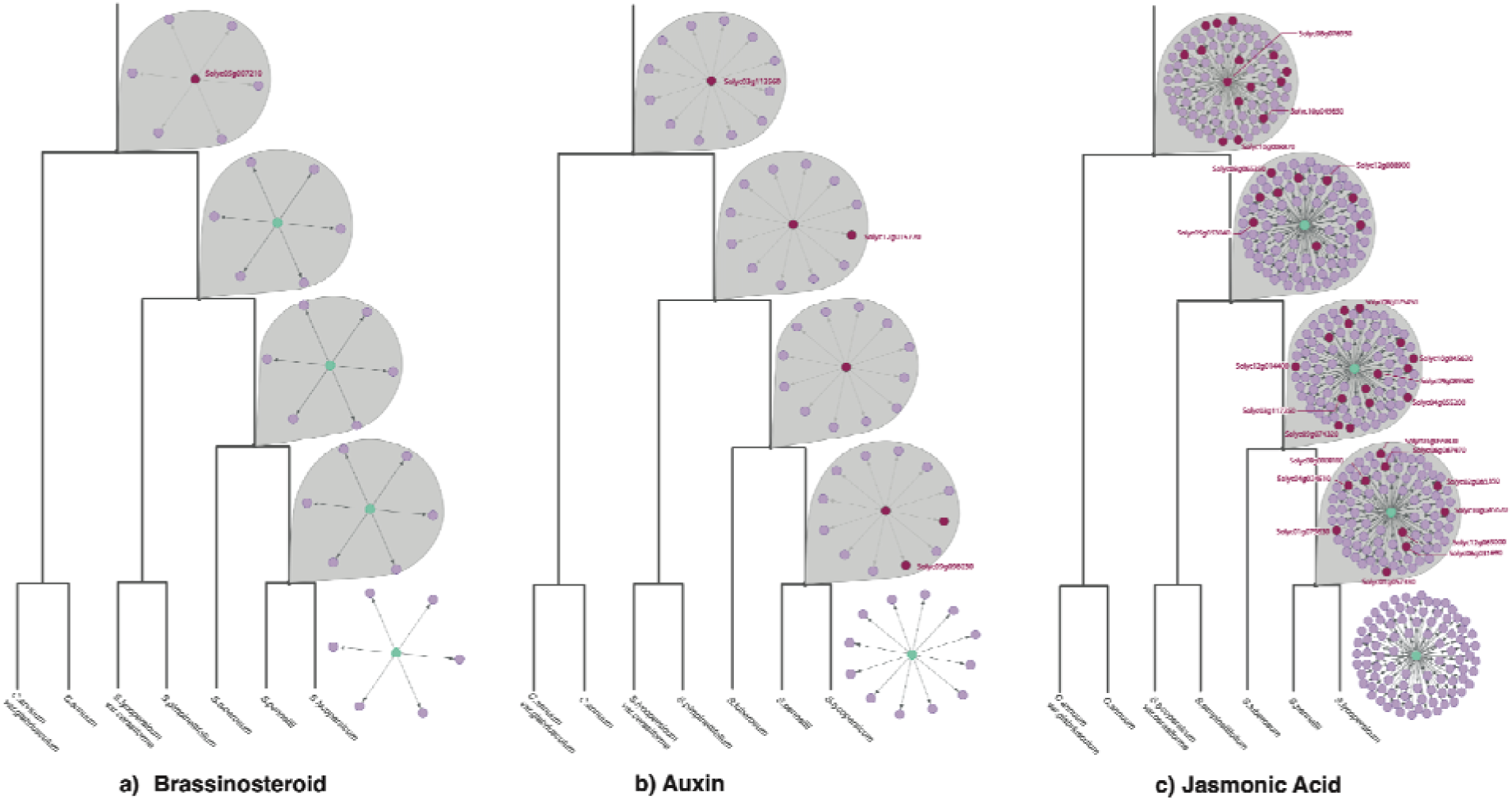
Conservation of regulatory and target genes in inferred networks associated with viroid-infection in tomato. a) In the regulatory network of Brassinosteroids during infection, the hub regulatory gene has orthologs in Solanum lineage but not in Capsicum. b) in Auxin inferred network, almost all target genes are conserved across species, however the bHLH regulatory gene is only present in tomato across Solanaceae. c) Similarly to Brassinosteroids, the bHLH regulatory hub gene is conserved across the Solanum lineage but absent in Capsicum. There are several tomato target genes that have no orthologs in other species. Green color node depicts the regulatory bhHLH gene. Nodes in pink are target genes which have orthologs in tomato, while red nodes represent genes that have no orthologs in tomato.

On the other hand, *Solyc03g113560*, mip-bHLH regulator, is a hub in the inferred auxin regulatory network, but has no orthologs in any other of the species in study, **Fig 5b**. However, this gene has an ortholog in *Arabidopsis thaliana* [34] but its absence in other Solanaceae (except domesticated tomato) suggests several evolutionary and functional scenarios that could explain this pattern. One of them is independent evolution in different lineages, since the gene may have evolved independently in Arabidopsis and tomato through convergent evolution, meaning both species developed a similar function but through different evolutionary events. The bHLH TF family is large and prone to duplications and subfunctionalization, meaning Arabidopsis and tomato could have developed similar regulators from different ancestral genes. Moreover, almost all target genes in this regulatory network have orthologs in the other species, suggesting rewiring under stress conditions.

Finally, the analysis of the JA-associated regulatory network, regulated by *Solyc08g076930*, reveals distinct patterns of conservation and divergence across *Solanaceae* species, **Fig 5c**. Our results highlight progressive regulatory divergence from *Capsicum* species to domesticated tomato (*S. lycopersicum*). At the top of the phylogeny, *C. annuum var. glabriusculum* and *C. annuum* exhibit a regulon, marked by gene abscense, including the MYC2/IA bHLH-MRT (*Solyc08g076930)*, and genes such as *Solyc10g049850*(TIP41-like protein), and *Solyc10g008870 (Transcription elongation factor 1)*. This suggests an ancestral divergence in JA signaling within the *Capsicum* lineage. Moving along the phylogeny, *S. lycopersicum var. cerasiforme* and *Solanum pimpinellifolium* retain a broader regulon than *Capsicum*, but notable abscenses, such as *Solyc08g065350 (Histone deacetylase 11)*, *Solyc05g053040 (RING finger family protein)*, and *Solyc12g088900*, indicate partial conservation of the ancestral JA network. Further along, *S. tuberosum* preserves most of the ancestral regulon, exhibiting fewer abscences, including *Solyc12g014400(Cell differentiation protein rcd1)*, *Solyc03g117250(Flotillin 1)*, and *Solyc09g074320 (Serine/threonine-protein phosphatase)*. The network of *S. pennelli* closely related when compared to that of domesticated *S. lycopersicum* has gene losses, including *Solyc08g000810*, *Solyc04g024510*, and *Solyc01g079830*.

Overall, these results demonstrate divergences of the JA regulon, evolving from a compact and highly pruned form in *Capsicum* to an expansive and complex network in domesticated *S. lycopersicum*. These patterns highlight the impact of evolutionary pressures, including domestication, on reshaping jasmonic acid regulatory networks within the *Solanaceae* family.

### 3.4 Network Rewiring Under Viroid Infection: Healthy vs. Infected Plants

In healthy plants, bHLH TFs exhibit a structured and modular regulatory architecture, characterized by cohesive functional interactions and tightly connected hubs [36]. This modular organization ensures precise control of key biological processes. However, under viroid infection, the regulatory network undergoes a drastic reorganization. The bHLH TFs display increased promiscuity, engaging with a broader range of partners but in a less specialized manner [37]. This loss of specificity results in aberrant connections that may contribute to disease-associated phenotypes.When comparing the regulons of our analyzed networks in healthy conditions, the bHLH TFs have a shift in their role, **Fig 6**. For instance the bhlh miP associated with brassionosteroid pathways has a totally different regulon. Likely the bhlh miP linked to auxin pathways is modulating the expression of a distintic set of genes. Finally, we observed that under healthy conditions, the bHLH-MTR MYC2/IA3 shares the regulation of its regulon with over a thousand TFs.

**Fig. 6.**
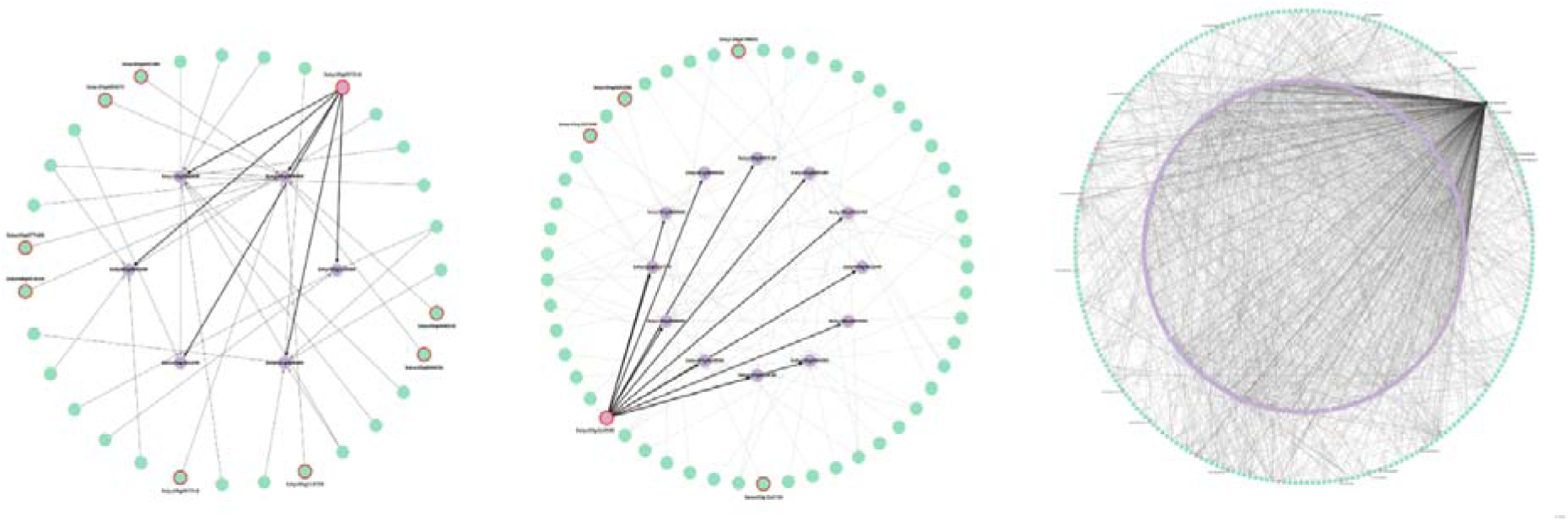
Rewiring of regulatory networks comparing viroid disease and healthy stages. Left: regulatory inferred network of the bhlh-mip brassinosteroid regulon during infection shows significant rewiring, with the bHLH regulating a set of genes that are regulated by other TFs in the healthy regulatory network. Middle: Auxin inferre regulatory network, similar to Brassinosteroids, has a bHLH hub gene targeting genes that are normally regulated by other TFs. Right: the inferred regulatory network of JA during infection shows a bHLH that regulated 91 genes that otherwise are regulated by more than thousand TFs in the healthy stage. Some of these TFs, bHLH as well, in the three networks, showing the highly specialized rewiring during pathogen intervention.

From an evolutionary perspective, transcription factor promiscuity is a key driver of regulatory plasticity and genetic innovation, allowing species to adapt to changing conditions through the rewiring of gene expression networks [35]. This reprogramming capability stems from the ability of bHLH TFs to form diverse homo- and heterodimeric complexes, enabling them to bind to a wide range of DNA sequences and regulate multiple gene expression pathways [36].

## 4. Discussion

Our study reconstructs the evolutionary trajectories of bHLH transcription factors (TFs) in Solanaceae and integrates this information with regulatory network analyses to understand their roles during PSTVd infection. By combining phylogenomic approaches with functional and network inferences, we reveal how evolutionary pressures, including domestication and viroid stress, have shaped bHLH-mediated regulatory mechanisms in tomato and its relatives.

The phylogenomic reconstruction highlights contrasting evolutionary trajectories between wild and domesticated Solanum species, shaped by gene gains, losses, and duplications. Notably, the domesticated tomato (*S. lycopersicum*) retains two species-specific bHLH genes (Solyc02g067380 and Solyc06g070910) with predicted roles in cytoskeletal organization and hormonal signaling. Their absence in wild relatives suggests that these genes may have emerged through domestication-driven selection, potentially contributing to traits such as growth regulation and stress adaptation. Conversely, the wild species *S. pennellii* shows a distinct evolutionary path, marked by multiple lineage-specific duplications. These duplications, which increase the number of co-orthologs, may represent adaptive responses to harsher environments, aligning with previous findings that associate gene duplication events with enhanced regulatory flexibility.

The inferred GRNs of *S. lycopersicum* under PSTVd infection reveal distinct regulatory modules for auxin, JA, and brassinosteroid pathways. Each module is governed by a key bHLH regulator, demonstrating functional specialization within the bHLH TF family. Our results support that bHLH-mediated hormonal crosstalk is central to coordinating transcriptional reprogramming during stress responses. The JA network, regulated by *Solyc08g076930* (IA3/MYC2), is highly conserved across Solanaceae species, highlighting the evolutionary importance of JA signaling in plant defense. Increased numbers of orthologs and duplications in potato (*S. tuberosum*) and wild tomatoes suggest that this regulatory module has been under strong selection for stress adaptation. In contrast, the auxin network, regulated by *Solyc03g113560* (miP-bHLH-ARF8), displays low evolutionary conservation, with the regulatory hub gene exclusive to domesticated tomato. Despite the conservation of target genes, the absence of this hub gene in other Solanaceae species indicates regulatory rewiring, likely driven by neofunctionalization following a duplication event. This divergence underscores its potential role in domestication-related traits, such as fruit development and stress adaptation. The brassinosteroid network, controlled by *Solyc05g007210* (miP-bHLH-BR), shows moderate conservation across Solanum species but a complete absence of the regulatory hub gene in Capsicum. This pattern indicates that the Solanum lineage may have evolved distinct brassinosteroid-mediated stress responses, potentially involving lipid metabolism and membrane remodeling, as suggested by the functional enrichment of target genes.

A comparison between healthy and PSTVd-infected GRNs reveals significant regulatory rewiring, characterized by changes in target gene connectivity and regulatory promiscuity. In particular, the JA-associated bHLH (*Solyc08g076930*) shifts from a co-regulatory architecture— where its regulon is controlled by over a thousand TFs in healthy conditions—to a dominant regulatory role under infection, where it directly governs 91 genes. This shift in regulatory control exemplifies the plasticity of bHLH TFs, allowing them to rapidly reconfigure transcriptional programs in response to biotic stress. Similar patterns of rewiring are observed in the auxin and brassinosteroid regulons, where bHLH regulators assume control over genes typically regulated by other TF families in healthy plants.

Our ortholog analysis reveals distinct patterns of evolutionary constraint and divergence among hormone-associated bHLH regulons. JA-associated bHLH regulons show strong evolutionary conservation, consistent with their well-established roles in stress defense. In contrast, auxin-associated regulons are more lineage-specific, indicating greater evolutionary plasticity, possibly due to their involvement in processes more closely linked to species-specific traits, such as development and reproduction. Brassinosteroid regulons display a balance between conservation and divergence, suggesting a dual role in both developmental and stress-adaptive pathways. The observed regulatory shifts under viroid stress highlight the ability of bHLH TFs to modulate plant defense networks through adaptive rewiring. This plasticity is likely a key factor in the evolutionary success of Solanaceae species in diverse and pathogen-rich environments. Additionally, our findings suggest that domestication may have inadvertently selected for bHLH-mediated traits that favor productivity and growth at the expense of regulatory diversity, as evidenced by the reduced complexity of JA-associated networks in domesticated tomato compared to its wild relatives.

## 5. Conclusion

Our study highlights the interplay between evolutionary conservation and regulatory rewiring in bHLH transcription factors of Solanaceae under domestication and viroid stress. The identification of lineage-specific bHLH regulators and their role in hormonal network modulation during PSTVd infection underscores their adaptive significance. Domestication has likely streamlined regulatory complexity, potentially impacting stress resilience. Future research should focus on functional validation of miP-bHLH regulators, their roles in viroid tolerance, and evolutionary analyses across Solanaceae, contributing to the development of resilient crop varieties through targeted gene editing strategies.

## Appendix

**Table 1.**
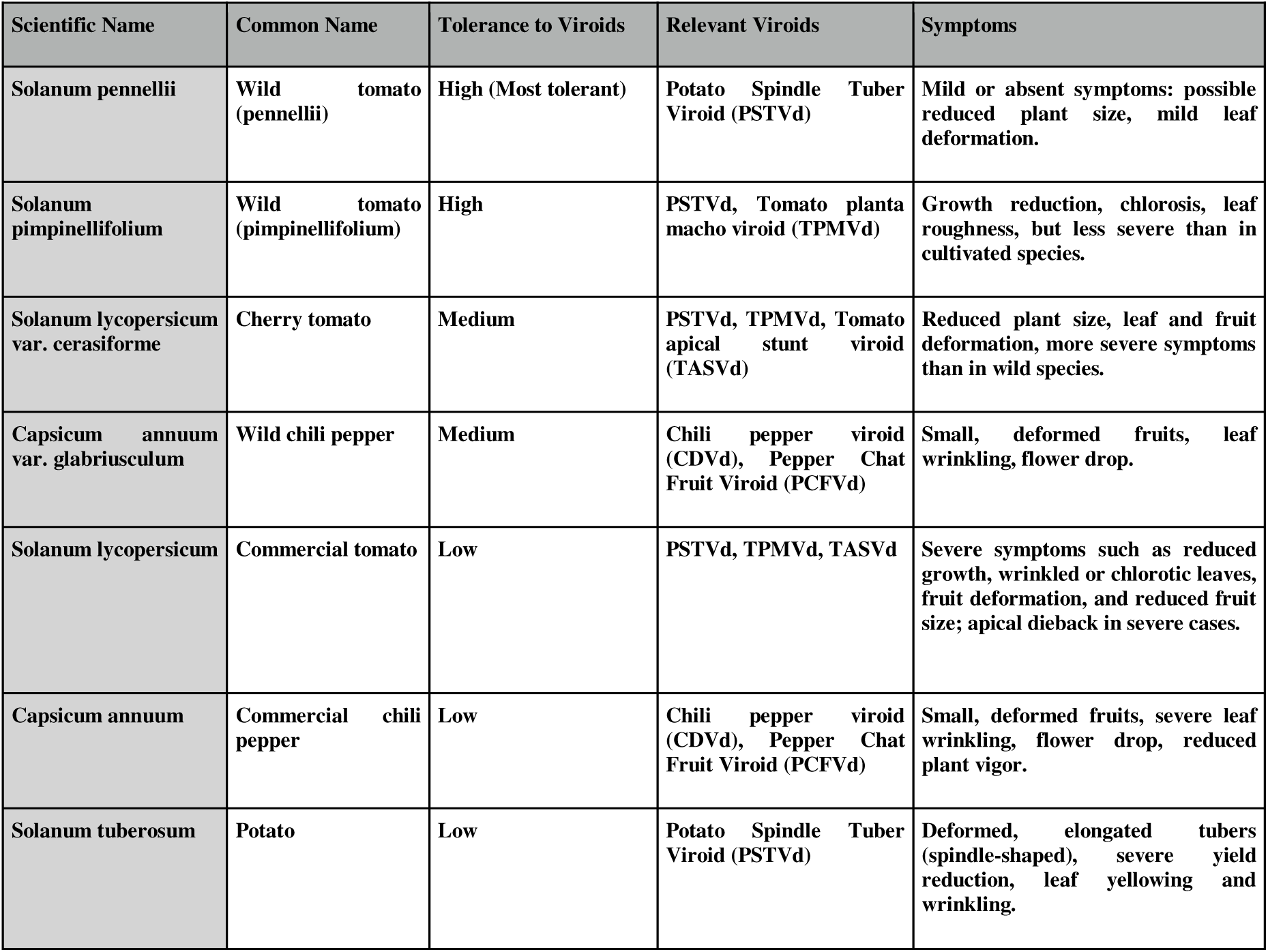
Relationship between plant species, associated viroids, and observed symptoms according to their tolerance level.

**Fig. A1.**
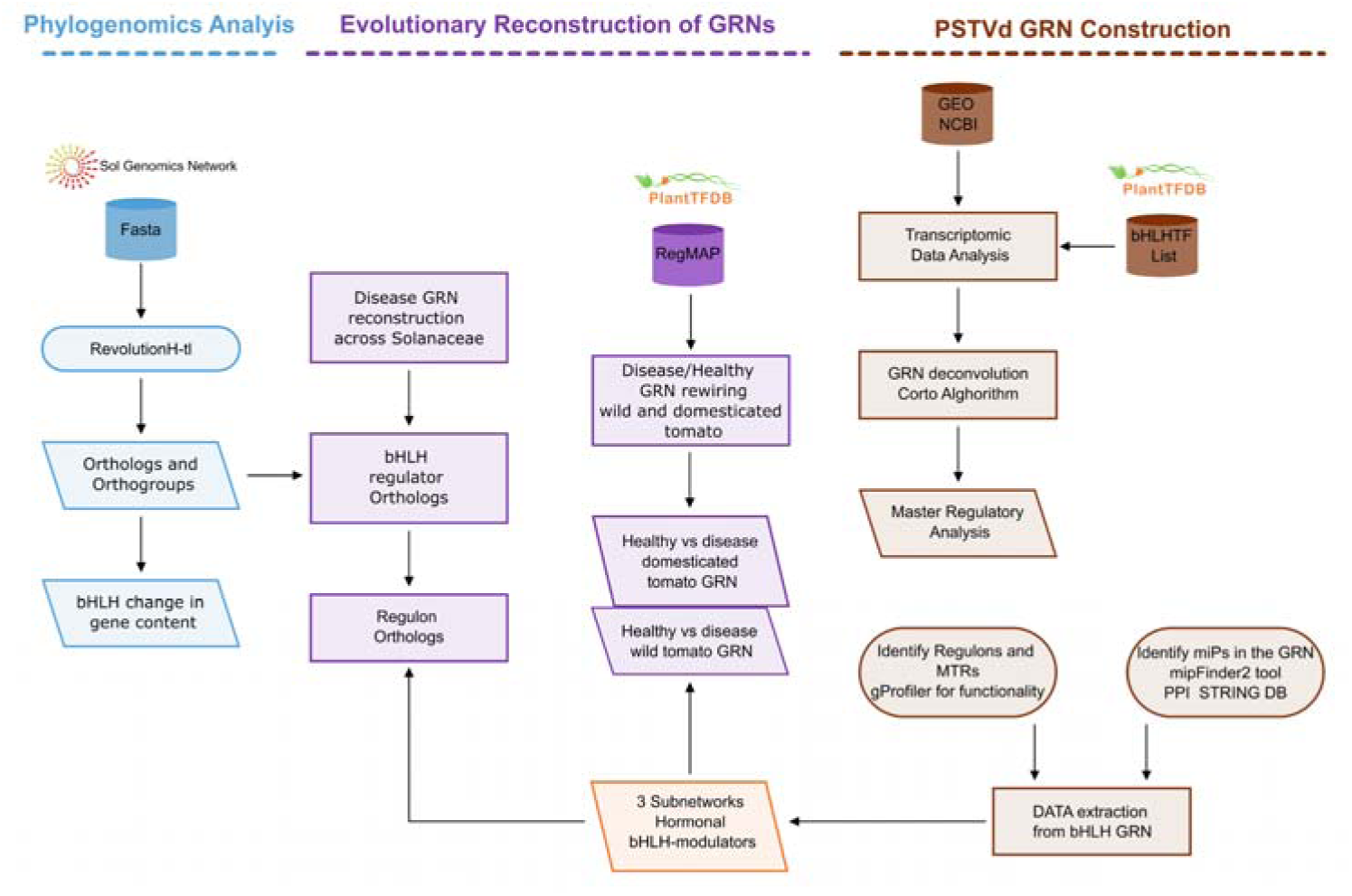
Workflow for Evolutionary Reconstruction and PSTVd-Induced GRN Analysis. The pipeline integrates three main approaches: (1) Phylogenomics Analysis to identify bHLH orthologs and gene content changes across *Solanaceae* using RevolutionH-tl. (2) Evolutionary Reconstruction of GRNs, comparing healthy and disease states in wild and domesticated tomatoes to identify bHLH regulons and hormonal subnetworks. (3) PSTVd GRN Construction, involving transcriptomic analysis, GRN deconvolution, and master regulatory analysis to identify bHLH targets and microproteins (miPs) using Corto and STRING databases.

**Fig. A2.**
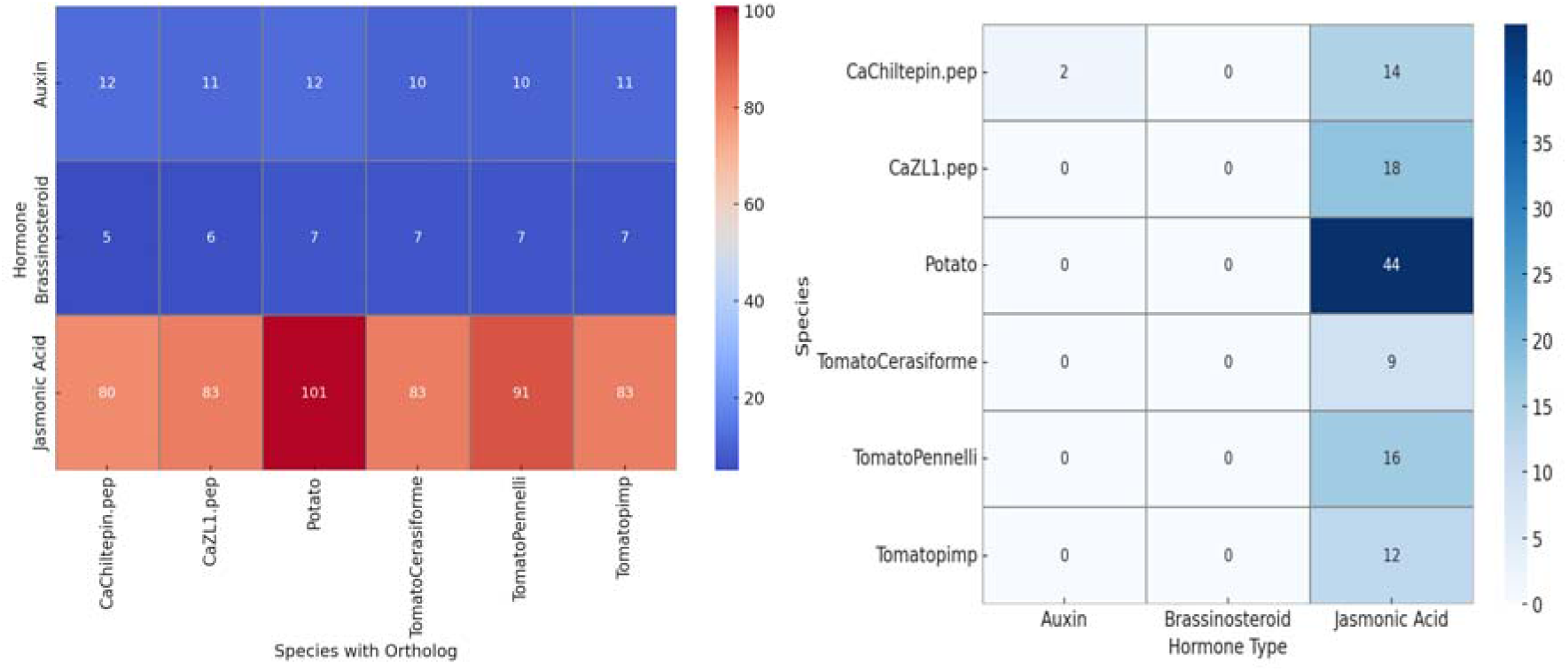
Heatmaps showing ortholog counts (left) and duplicated orthologs (right) for bHLH-ARF8, bHLH-brassinosteroid and IA3/MYC2 regulons across Solanaceae. JA genes are the most abundant and duplicated, with potato showing the highest duplications. Auxin and brassinosteroid regulons display lower counts and duplications, indicating distinct evolutionary patterns.

## Notes

### Competing Interest Statement

The authors have declared no competing interest.

