## Appendix for "Evolutionary Reconstruction of Hormone-bHLH Regulatory Networks in Solanaceae: Phylogenomic Insights from PSTVd-Tomato Interactions"

**Table 1. Relationship between plant species, associated viroids, and observed symptoms according to their tolerance level.**

| **Scientific Name** | **Common Name** | **Tolerance to Viroids** | **Relevant Viroids** | **Symptoms** |
| --- | --- | --- | --- | --- |
| **Solanum pennellii** | **Wild tomato (pennellii)** | **High (Most tolerant)** | **Potato Spindle Tuber Viroid (PSTVd)** | **Mild or absent symptoms: possible reduced plant size, mild leaf deformation.** |
| **Solanum pimpinellifolium** | **Wild tomato (pimpinellifolium)** | **High** | **PSTVd, Tomato planta macho viroid (TPMVd)** | **Growth reduction, chlorosis, leaf roughness, but less severe than in cultivated species.** |
| **Solanum lycopersicum var. cerasiforme** | **Cherry tomato** | **Medium** | **PSTVd, TPMVd, Tomato apical stunt viroid (TASVd)** | **Reduced plant size, leaf and fruit deformation, more severe symptoms than in wild species.** |
| **Capsicum annuum var. glabriusculum** | **Wild chili pepper** | **Medium** | **Chili pepper viroid (CDVd), Pepper Chat Fruit Viroid (PCFVd)** | **Small, deformed fruits, leaf wrinkling, flower drop.** |
| **Solanum lycopersicum** | **Commercial tomato** | **Low** | **PSTVd, TPMVd, TASVd** | **Severe symptoms such as reduced growth, wrinkled or chlorotic leaves, fruit deformation, and reduced fruit size; apical dieback in severe cases.** |
| **Capsicum annuum** | **Commercial chili pepper** | **Low** | **Chili pepper viroid (CDVd), Pepper Chat Fruit Viroid (PCFVd)** | **Small, deformed fruits, severe leaf wrinkling, flower drop, reduced plant vigor.** |
| **Solanum tuberosum** | **Potato** | **Low** | **Potato Spindle Tuber Viroid (PSTVd)** | **Deformed, elongated tubers (spindle-shaped), severe yield reduction, leaf yellowing and wrinkling.** |

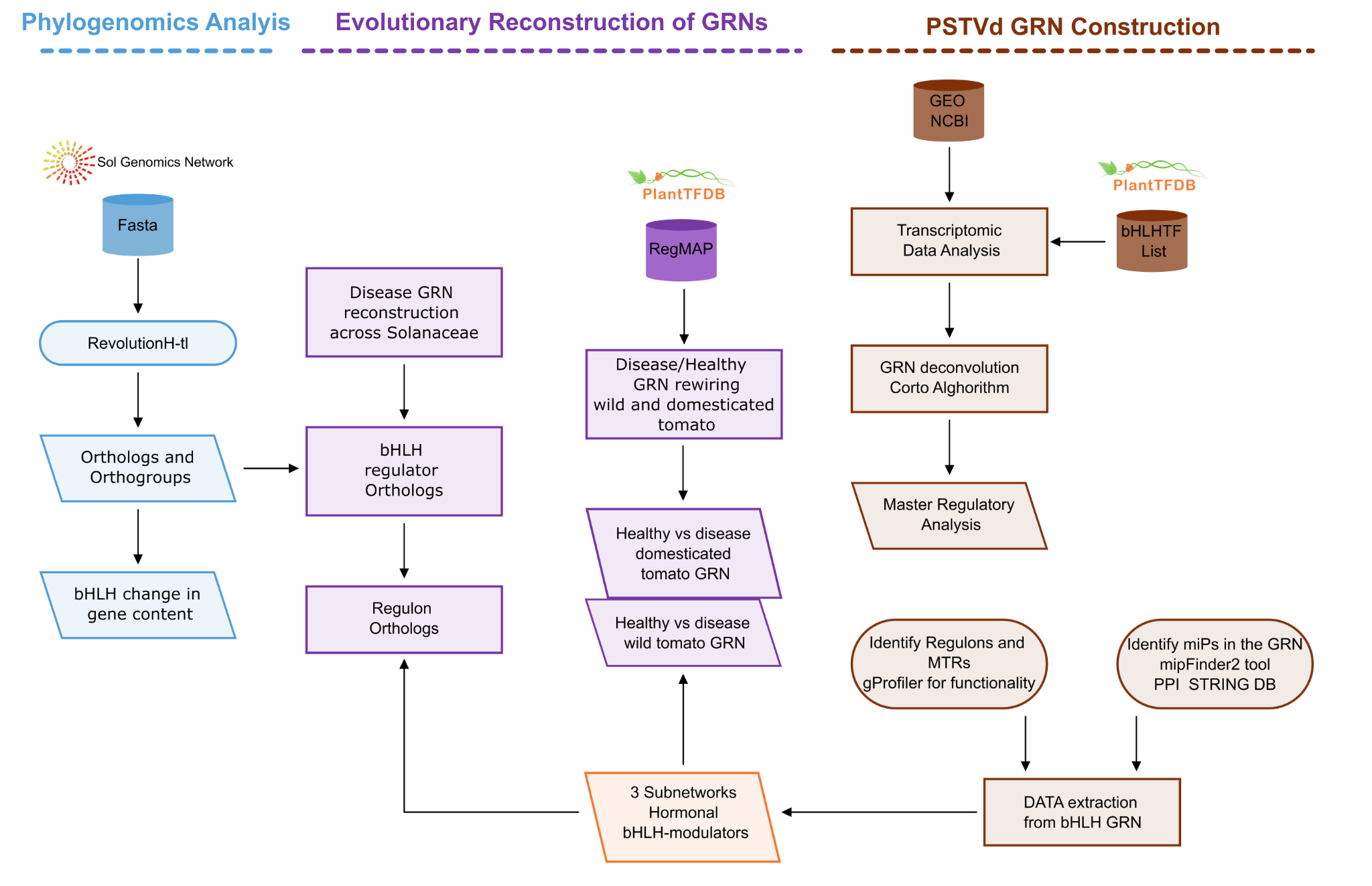

**Fig. A1.Workflow for Evolutionary Reconstruction and PSTVd-Induced GRN Analysis.** The pipeline integrates three main approaches: (1) Phylogenomics Analysis to identify bHLH orthologs and gene content changes across *Solanaceae* using RevolutionH-tl. (2) Evolutionary Reconstruction of GRNs, comparing healthy and disease states in wild and domesticated tomatoes to identify bHLH regulons and hormonal subnetworks. (3) PSTVd GRN Construction, involving transcriptomic analysis, GRN deconvolution, and master regulatory analysis to identify bHLH targets and microproteins (miPs) using Corto and STRING databases.

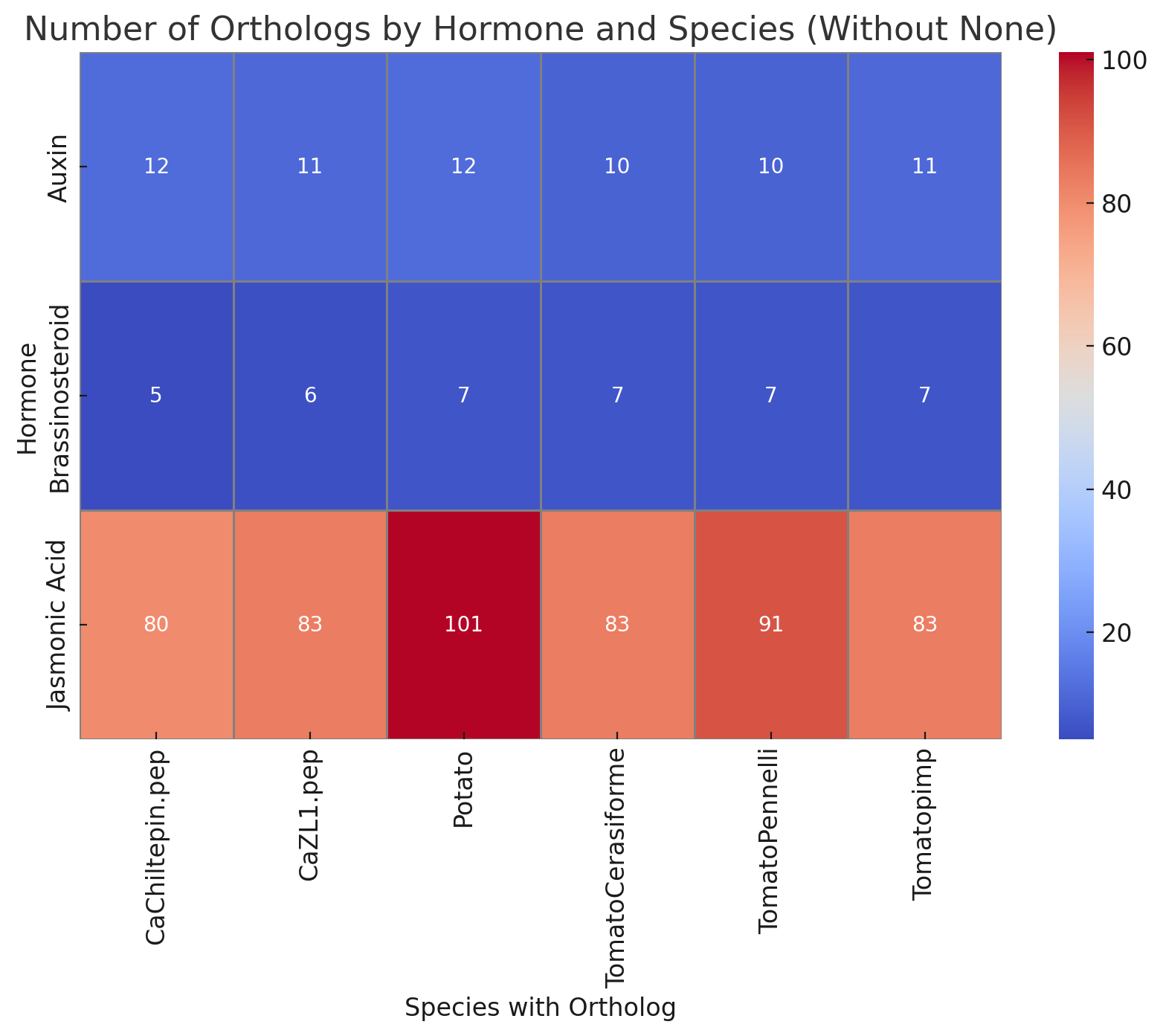

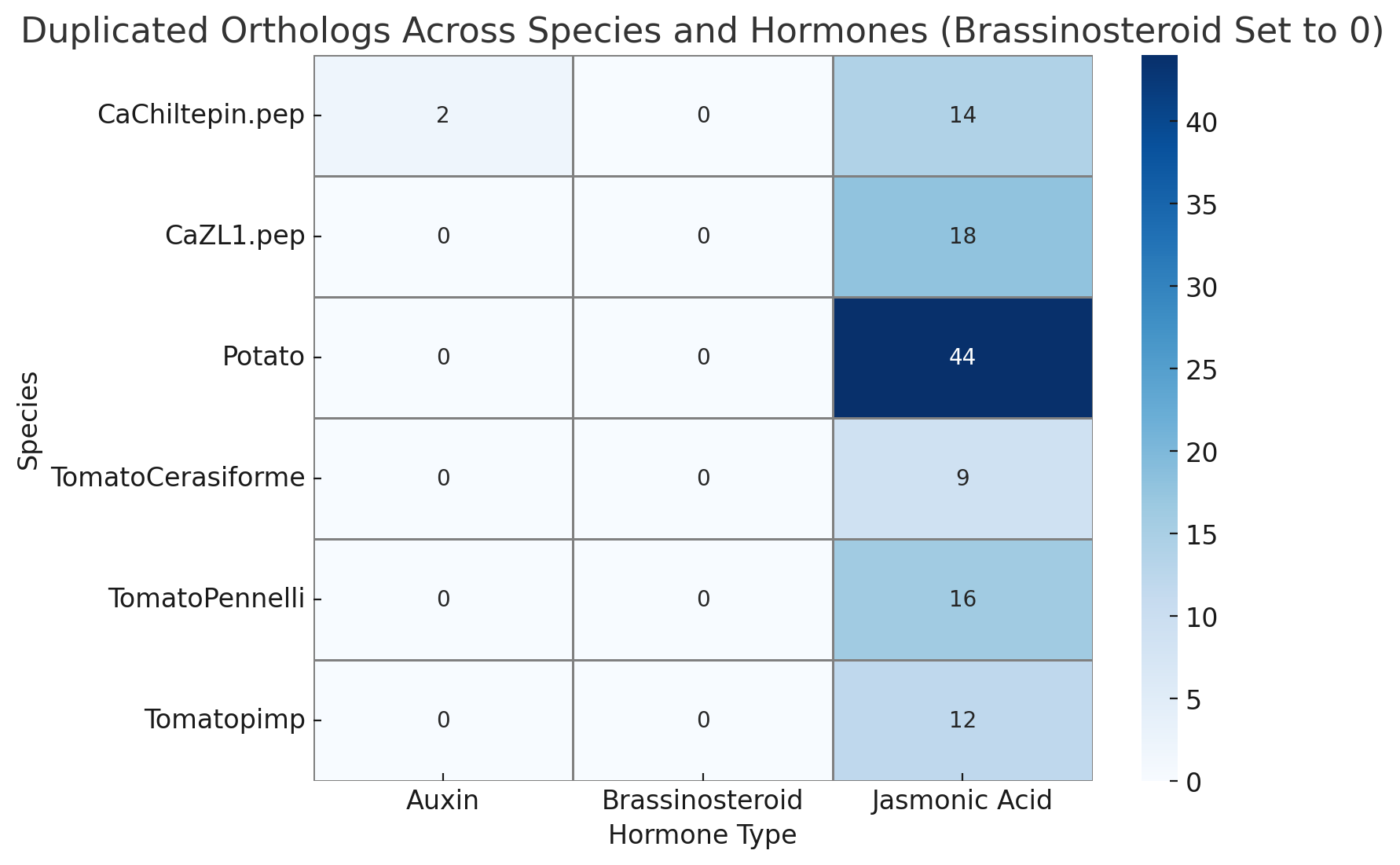

**Fig. A2.** **Heatmaps showing ortholog counts (left) and duplicated orthologs (right) for bHLH-ARF8, bHLH-brassinosteroid and IA3/MYC2 regulons across Solanaceae.** JA genes are the most abundant and duplicated, with potato showing the highest duplications. Auxin and brassinosteroid regulons display lower counts and duplications, indicating distinct evolutionary patterns.
